# Home is where the host is: Evolutionary history of geographic spread, host switching, and adaptive genomic signatures in two generalist Group B *Streptococcus* clonal groups

**DOI:** 10.64898/2026.01.06.697867

**Authors:** Antonia Hilbig, Chiara Crestani, Timothy Barkham, Swaine L. Chen, Claudia Cobo-Angel, Alejandro Ceballos-Marquez, Wanna Sirimanapong, Syafinaz Amin-Nordin, Nguyen Ngoc Phuoc, Tatiana Castro Abreu Pinto, Laura Oliveira, Chiara Anna Garbarino, Matteo Ricchi, Tiziana Lembo, Samantha Lycett, Roman Biek, Taya Forde, Ruth N. Zadoks

**Author notes:** These authors contributed equally (Roman Biek, Taya Forde, Ruth N. Zadoks).

## Abstract

Group B *Streptococcus* (GBS) is a pathogen of global relevance in neonatal and maternal disease as well as bovine mastitis. Two closely related clonal groups, denoted 103 and 314 (CG103/314) have been detected in humans and cattle on multiple continents in recent decades but are poorly characterised compared to other host-generalist clades. We examined their potential origins, host-switching events and presence of a suite of genetic markers for antimicrobial resistance, virulence and host association using a newly assembled dataset of 248 CG103/314 genomes from humans, cattle, and food originating from five continents. We detected multiple host switches between humans and cattle, and significant regional differences in AMR gene distribution, possibly reflecting local differences in antimicrobial use across countries and hosts and indicating a capacity for regional adaptation to selective pressures. Across the evolutionary history of CG103/314 from both host species, the prevalence of the Lac.2 operon, a genetic marker associated with bovine host adaptation, was high, whereas the prevalence of the *scpB-lmb* gene pair, a genetic marker of human host adaptation in other GBS clonal groups, was very low. All isolates with *scpB-lmb* were associated with human disease rather than carriage. Our dataset displayed biases typical of research into multi-host pathogens, when sampling is often focused on a specific host species or setting. Consistent, balanced, contemporaneous and sympatric sampling efforts across host species and sources are needed for a full understanding of the distribution and emergence of CG103/314 and similar multi-host pathogens impacting food safety and public health.

**Impact statement:** This study provides a comprehensive, global genomic overview of generalist clonal groups 103/314 of the human and animal pathogen Group B *Streptococcus* (GBS). By analysing host switching, antimicrobial resistance and virulence-associated markers, we show that these clonal groups display adaptation patterns shaped by region- and host-specific selective pressures. Our findings include potential expansion of the host range from humans and cattle into porcupines and pigs, and provides detailed discussion around anthropocentric sampling bias, highlighting the importance of balanced, multi-host sampling of generalist GBS lineages and One Health pathogens in general. This work reinforces the need for coordinated One Health surveillance to monitor emerging sub-lineages with relevance for food safety, human and animal health.

## Introduction

*Streptococcus agalactiae*, also known as Group B *Streptococcus* (GBS), is a gram-positive bacterium initially discovered as a cause of bovine mastitis, hence named ’a-galactiae’ meaning ’without milk’ (1–3). GBS has a broad host range, colonising or infecting humans (4), cattle (5), camels (6) and teleost fishes (7), and occasionally other species as taxonomically diverse as dogs and cats (8), sea mammals, reptiles, amphibians, and cartilaginous fishes (9, 10). In humans, GBS is notorious for causing severe neonatal disease and stillbirths, with approximately 160,000 - 230,000 cases of invasive disease in infants each year globally (11) and levels of maternal colonisation averaging 18%, with significant local heterogeneity (12, 13). The average prevalence of GBS colonisation is similar in non-pregnant adults (14), and invasive GBS disease is increasingly recognised in adults, particularly in elderly patients with comorbidities (15) and, in Asia, due to the fish-associated strain ST283 (16, 17). In cattle, GBS causes bovine mastitis, leading to significant economic losses in the dairy industry (18).

A recent study has identified host specialist and host generalist lineages within GBS (19). Generalist pathogens are of concern to human and animal health, as targeting interventions towards multiple hosts is challenging and host generalists may persist within non-targeted environments or hosts, creating an opportunity for re-emergence. An example is the emergence of sequence type (ST) 103 and ST314 in Swedish cattle after a period when GBS seemed to have been eliminated from the dairy cattle population, with humans postulated to be the source of re-emergent GBS in cows (20). Sequence type 103 and its single locus variant ST314 are the namesake ST of clonal groups (CG) 103 and 314, which form a monophyletic clade within the global GBS population (supplementary Figure S1). A non-bovine evolutionary origin for mastitis-causing CG103/314 was suspected, in part because the stem group of this clade (ST314) carried a tetracycline resistance gene, despite tetracycline use being exceedingly rarely in Swedish dairy cattle (20). At the time, tetracycline resistance had been described as a marker of human host adaptation in GBS (21). Another unusual feature of CG103/314 in cattle is its detection in faecal and environmental samples, which does not match the historical veterinary paradigm that bovine GBS is an “obligate intramammary pathogen” (22, 23). As a result, standard measures that are recommended globally for control of GBS mastitis, and which focus on prevention of cow-to-cow transmission only, may not be adequate for control of CG103/314 (22, 24). Whether the unusual epidemiology, and the resulting failure of routine mastitis control practices, reflect a non-bovine origin of CG103/314 is speculative, because the historical association of tetracycline resistance with human GBS, which underpinned the idea of a human origin for bovine CG103/314, was not supported by subsequent genome wide association studies that consider GBS from a wide range of hosts and countries (19).

Both CG103 and CG314 were initially considered host specialists, as almost all available isolates originated from bovine milk based on studies in Europe (25, 26) and China (27). Later, more data from a wider diversity of locations, sources and timepoints became available and analyses focused on host-association. This altered our perception of these CGs to that of a host generalist (defined as a CG of which less than 80% of isolates originate from a single host species) (19). Human carriage has been reported from Brazil (28, 29) and severe cases of human disease have been reported from Asia (30–32). Detection of CG103/314 in bovine milk, either from the bulk tank or from cases of bovine mastitis, is common across multiple continents, including Asia (33), Europe (34, 35) and South America (36), with possible bi-directional transmission between people and cattle on dairy farms (34, 37). More recently, CG103/314 isolates have occasionally been reported as a cause of disease in pigs and wildlife (38) and in pigs’ organs collected from wet markets (39), suggesting that this CG may be expanding beyond its currently-recognised human and bovine hosts.

To explore the evolutionary dynamics and host-switching potential of CG103/314, we analysed a collection of publicly available and newly sequenced GBS genomes representing human, bovine and food surveillance isolates across multiple continents. In our analyses, we assessed i) AMR and virulence gene presence across hosts, location and ST, ii) then clarified population structure by use of phylogenetic and network analyses and iii) quantified host switching. Our characterisation of these CGs provides a comprehensive foundation for future investigations into this global One Health pathogen.

## Methods

### Dataset curation

A literature search was conducted to identify publications (up to and including 2023) analysing GBS isolate collections that contained at least one CG103/314 isolate. Search terms used at PubMed/NCBI were combinations of “GBS”, “Group B *Streptococcus*”, “*Streptococcus agalactiae*”, “*Streptococcus*” in combination (search syntax “AND”) with “103”, “314” and “651” each preceded by respectively “ST”, “CC”, “CG” or “sequence type”, “clonal complex” or “clonal group” or the numbers by themselves. We additionally scanned literature cited in the publications that were retrieved in this way. In total we identified 42 publications and 2 non-published collections (personal communication M. Bijlsma, Amsterdam University Medical Center, The Netherlands; and unpublished observations J. F. Bohnsack as mentioned in Oliveira et al. (2006)) that include at least one CG103/314 isolate (40). One publication was excluded from subsequent analysis as there was insufficient information on the number of ST103 isolates (41). Figure 1 presents a visualisation of all isolates and isolate collections considered in this study. These include both isolates for which sequence data were not available, as well as isolates for which sequence data were available and analysed in the following sections.

**Figure 1.**
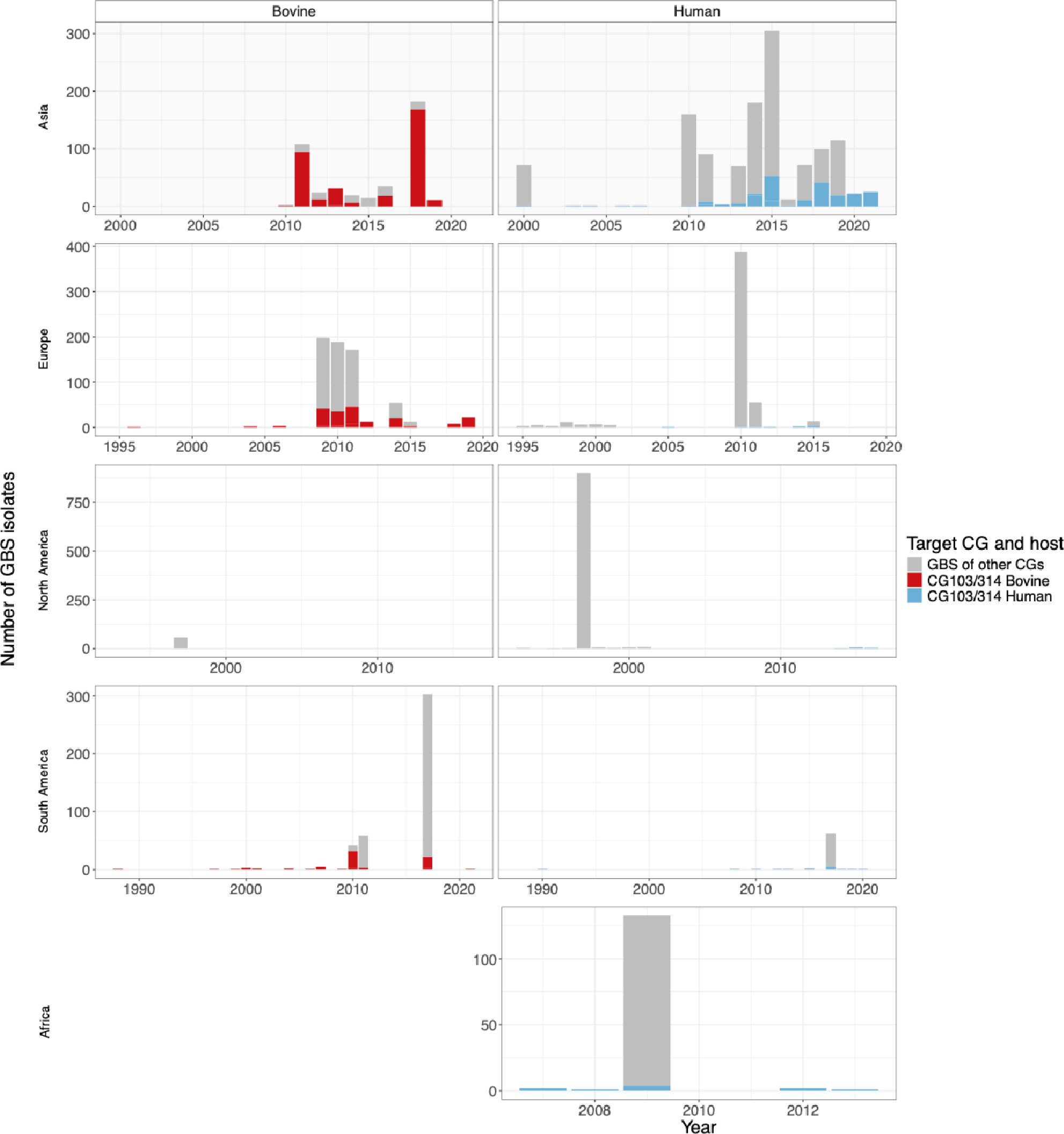
Distribution of clonal group (CG) 103/314 and non-CG 103/314 Group B Streptococcus isolates in collections that include at least one member of clonal group (CG) 103 or 314. Grey bars represent the number of GBS isolates reported per year and region, with coloured segments representing human (blue) or bovine (red) CG103/314 isolates. Three isolate collections from Western Asia (Israel) and Oceania (New Zealand) were excluded from visualisation due to insufficient data points. When time ranges for sample collection were given instead of specific years, only those collections with a collection interval no greater than 5 years were included in this visualisation and the midpoint of the range was chosen for representation (see Table S1 for a full list of publications and exact time ranges of all isolates from collections).

Scientific literature pre-dating 2024 documents the presence of CG103/314 in both humans and bovines, and a scarcity of data from continents other than Asia, Europe and South America despite an increasing number of detections globally after 2010 (Figure 1). Both CGs are reported from both host species in the three named continents, with an increasing number of detections in humans after the middle of the 2000s. In the late 2000s and middle of the 2010s (42–44), CG103/314 has occasionally been reported from humans in North America, whereas the only collection containing CG103/314 isolates from Africa was published in 2011 (45).

We went on to identify isolates that had been subjected to whole genome sequencing by conducting a comprehensive search of the above-mentioned isolate collections and through the research activities of the authors. An overview of isolates with whole genomes available versus those without is presented in the supplementary Figure S2.

Genomes were included in our analyses if they belonged to ST103, ST314 or their single, dual or triple locus variants as defined by multilocus sequence typing (MLST) of seven housekeeping genes (46). Genome sequences comprised those obtained from public databases (accession numbers are provided in the Microreact Project, a link is provided at the end of this article) and isolates that were sequenced for this study. Ten bovine isolates representing ten farms in Italy and known to belong to ST103 based on amplicon sequencing, as well as one isolate each from a pig and a porcupine (38) were submitted for whole genome sequencing by MicrobesNG (Birmingham, UK) using the Illumina platform. Isolates from Singapore, Malaysia, Thailand and Vietnam were sequenced as described in Sirimanapong et al. (2023) using the Nextera library preparation system and Illumina platform (47).

### Sequence processing and quality control

Of the 248 genomes, 91 were retrieved from public databases and 157 had been generated by the authors as part of their ongoing research into GBS epidemiology (and were made public after dataset curation). Nineteen genomes were available as assemblies and the remaining 229 were available as raw reads (see associated Microreact project for all accession numbers).

The fq2dna pipeline (https://research.pasteur.fr/en/tool/fq2dna/) was used to trim and assemble raw reads *de novo*. As fq2dna can only process paired-end read data, three single-end libraries were assembled into contigs using SKESA 2.5.1 (48). Genomes were annotated using Prokka (49). Quality control of genome assemblies was carried out using QUAST 5.2.0 (50) and results for the total length of the genome, total number of contigs, N50, L50 and GC content were plotted using the ggplot2 package in R (51) (see Figure S3). Genomes with values of these metrics exceeding twice the standard deviation of the data distribution’s mean were examined separately with KmerFinder 3.0.2 (Database version: 2022/07/11) with a query against all bacterial organisms (52–54).

### Descriptive genomic analysis

Commonly used typing systems for GBS are capsular serotyping (CPS), MLST and clustering algorithms. Capsular serotypes were determined using the GBS-SBG script and reference library (55), which identifies all ten serotypes of GBS (Ia, Ib, and II through IX) as well as subserotypes III-1 through III-4. Sequence types were determined using PubMLST (56, 57). There were 16 genomes for which we could not assign an ST; these were denoted as non typable (“NT”, see Microreact Project). Of these, 10 had an atypical *glcK* allele (58) and the remainder had new alleles or allelic profiles and were submitted to PubMLST for ST assignment (outcome pending).

Genomes were screened for the Lac.2 operon and the genes *scpB* and *lmb* using blastp (59) on amino-acid databases for each of the gene sets (see also supplementary material section 1). The genomes were assessed for the presence of antimicrobial resistance (AMR) genes against three different databases, namely ResFinder (60), the Comprehensive Antimicrobial Resistance Database (CARD) (61) and MEGARES v. 3.0.0 (62), using abricate (https://github.com/tseemann/abricate). The latter was also used with the virulence factor database VFDB (63) to detect virulence genes.

The association of the presence of *scpB-lmb* with human disease (as opposed to carriage) was tested using Fisher’s exact test for a subset of genomes where information on human disease status was available (95 out of 130 genomes with human origin had information on any disease. For information on disease status, please see Microreact Project).

### Genome alignment, network analysis and phylogeny construction

Using snippy (https://github.com/tseemann/snippy), a whole genome SNP alignment was generated, containing those mono- and polymorphic sites present in parts of the genome conserved among all genomes included in the analysis (core sites), using the largest genome in the dataset as reference (SRR6327910). For unresolved relationships between genomes that could be caused by recombination in the core genes, SplitsTree v4.19.2 (64) was employed to generate a Network with the NeighbourNet method (65) using the whole genome SNP alignment not filtered for recombination. Subsequently, Gubbins v3.1.0 (66) was used to detect and remove recombination. Recombination blocks were visualised in Phandango (67) together with a maximum likelihood phylogeny for the full dataset of 248 genomes, inferred with IQ-Tree version v2.2.2.3 (68) (see Figure S4).

Network analysis and phylogenies generated with maximum likelihood methods use sequence data only, whereas the BEAST package additionally uses time-stamp data to infer time-scaled trees, assuming a molecular-clock model applies (69). TempEst was used to assess clock-like behaviour among all genomes with an available year of isolation (n = 246, no year information on SRR494339 and CP010319) (70). We used a maximum likelihood phylogeny and performed temporal rooting along with root-to-tip regression using the correlation algorithm (“find best fitting root” option). Outliers were assessed and one sequence (GBS from a milk sample, ERR2729325) was excluded because of a strong mismatch between phylogenetic placement and year-information. A positive slope in the root to tip regression confirmed a clock-like behaviour (see Figure S5). This left a recombination-filtered core SNP alignment with 245 sequences and 2717 sites for time-scaled analyses using BEAST. A list of these 245 genomes with their metadata is available in the Microreact Project. For ancestral state reconstruction analysis (see next section), we excluded genomes from food isolates, pig and porcupine, leaving an alignment of 205 sequences from human or bovine hosts.

### Phylodynamic analyses

We inferred ancestral states in the phylogenies of CG103/314 using metadata on the host of origin of sequenced isolates and two Bayesian phylodynamic approaches: discrete trait (DT) mapping as implemented in BEAST-1 (71, 72) and the structured coalescent as implemented in the MASCOT package v3.0 in BEAST-2 (73–75). We also used DT to infer the ancestral presence of genes that were identified as markers of host-association in genome wide association studies, i.e., *scpB-lmb* and the Lac.2 operon (19).

To explore the potential impact of sampling bias, DT analysis was also conducted using continent-host combinations as ancestral state (e.g., European bovine, South American human) and using subsampled datasets for host or continent-host combinations. These are described in the supplementary material.

Time-scaled phylogenetic trees were inferred using BEAST v1.10.4 (69, 76). Initial model selection was performed by comparing the fit of substitution models, clock models and effective population size tree priors, as assessed by posterior parameter distributions in Tracer (77), using marginalised likelihoods based on Path sampling and Stepping Stone sampling (78, 79). Final models were run using a strict molecular clock, Bayesian Skyline population model and a general time-reversible (GTR) substitution model. Models were fitted in three independent runs with a Markov chain Monte Carlo (MCMC) chain of length 200,000,000, sampling every 20,000 iterations, reaching an effective sampling size of >300 for all parameters and discarding the first 10% of samples as burn-in. To ensure that tree topology was informed solely by the sequence and sampling date information, a posterior set of 1000 trees was subsampled from an analysis run without traits. Presence of host-associated marker genes and continent-host combinations were reconstructed upon these trees in BEAST using asymmetric DT models with Markov Jumps and using Bayesian stochastic search variables selection (71). As Bayesian stochastic search variables selection allows for migration rates to be switched off during sampling phases (71), only the mean of all samples during which the respective rates were switched on was extracted manually from the log files. Visual representation of migration between the continents was produced using Cytoscape (80).

In the structured coalescent approach, (75)final models were run using a strict molecular clock, MASCOT population model and GTR substitution model. The effective population size prior was drawn from a uniform distribution to allow for flexibility in values as there are no pre-existing estimates in the literature for phylogenetic analysis of CG103/314 to base assumptions on. The constant migration rate priors were drawn from a lognormal distribution with mean in real space of 0.002 and a standard deviation of 1.5, favouring small values without excluding large values, based on exploratory analyses in BEAST-1. An exponential distribution was rejected after exploratory analyses in MASCOT. Xml-files were manually modified to allow for asymmetric migration between the host or continent-host demes.

Models were fitted in three independent runs with an MCMC chain of length 200,000,000 sampling every 20,000 reaching an effective sampling size of >1000 for all estimates and discarding the first 10% of samples as burn-in. We reconstructed CG103/314’s demographic history using a Bayesian Skyline prior (Figure S6).

## Results

### Data Summary

All sequence data used in this study is publicly available and accession numbers can be viewed and downloaded from the metadata of the Microreact project; URL: https://microreact.org/project/qdpMsBtAX8FJmTXVZm85AZ-gbscg103314hilbigetal

### Dataset

The final dataset of 248 CG103/314 genomes originated from GBS that were isolated from 18 countries and five continents from 1990 to 2023 (inclusive), predominantly from humans (n = 130), cattle (n = 78) and food (n = 38) (Figure 2). Food samples were either raw fish or meat from food markets in Singapore (Figure S7). Wet market isolates from Hong Kong have been described (39) but sequence data were not available from the associated NCBI BioProject (PRJNA752017) before our inclusion cut-off date of Dec 31^st^ 2023.

**Figure 2.**
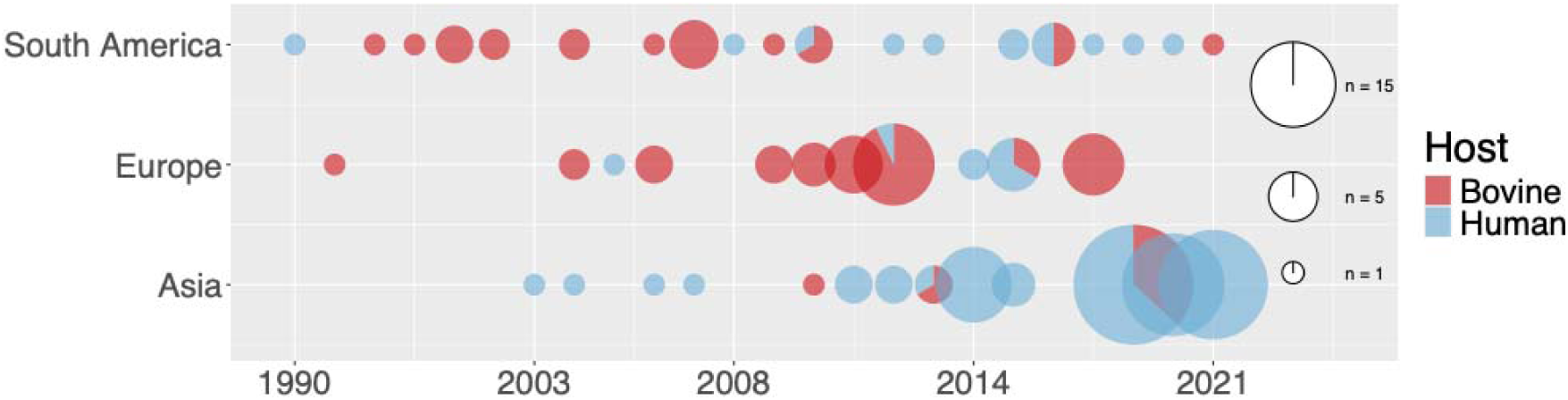
Geographical and temporal origin of the Group B Streptococcus clonal complex (CG)103/314 genomes in the dataset. Pie sizes represent the number of CG103/314 genomes with pie proportions and colours indicating the host of origin (human in blue, bovine in red). Genomes from Africa (n =6) and North America (n = 15) were excluded from the visualisation because of insufficient data. For a representation including food isolates, see supplementary Figure S7.

**Figure 3.**
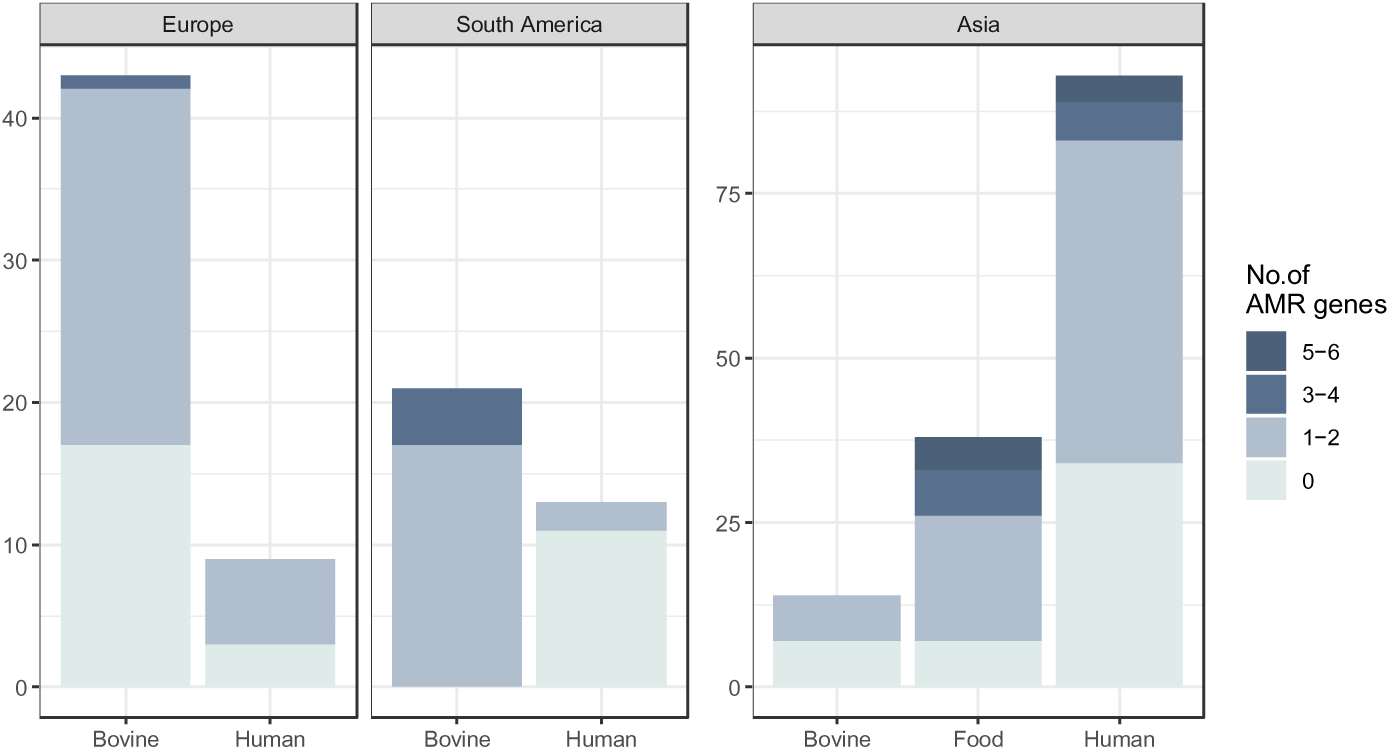
Prevalence of antimicrobial resistance (AMR) genes in genomes of Group B Streptococcus (GBS) belonging to clonal groups (CG) 103 and 314. Results are stratified by continent and sample type. Intensity of shading indicates the number of AMR genes. Results for Africa and North America are excluded because of an insufficient number of genomes. Note the difference in scale between results for Asia and those for other continents.

The dataset, which represents a convenience sample based on availability rather than purposive sampling, has three limitations. First, there is an imbalance with 60% of GBS genomes originating from human isolates and ∼ 30% from bovine isolates. This does not reflect the distribution of isolates described in the literature as depicted in Figure 1, where CG103/314 were proportionally more common among bovine than human GBS.

Secondly, the early years of the sampling interval are represented predominantly by bovine isolates. The one notable exception is the earliest genome, which originated from a human isolate collected in 1990 (Figure 2). These two features of the dataset likely have a strong yet competing effect on the inference of early host association. A third limitation is geographic, in that the majority of the human GBS genomes are of Asian origin, while the majority of the bovine GBS genomes are from Europe or South America. When comparing the traits of the sequenced isolates with the traits of non-sequenced isolates from the literature (Figure S2), it becomes apparent that there is a large population of Asian bovine associated CG103/314 strains that are not captured in our genomic analyses due to unavailability of genomic data.

However, the host distribution of CG103/314 genomes from Europe and South America broadly mirrors the proportions of isolates for those continents (Figures 1 and 2).

### Descriptive genomic analysis

Genome sizes ranged from 1,933,112 base pairs (bp) to 2,317,504 bp with a median of 2,070,054 bp and a mean of 2,084,208 bp. Genome size did not correlate with host origin. Median N50 and median number of contigs were 488415.5 and 15, respectively (for genome sizes, N50 values and numbers of contigs, see Microreact project).

Four CPS and sixteen ST were detected. Most isolates belonged to CPS type Ia (219 of 248 or 88%), with the remainder belonging to type III-3, VII and II (n = 21, n = 2 and n = 1 isolates, respectively). For five isolates, CPS could not be determined (Microreact project). The most common ST was ST103 (n = 134), followed by ST314 (n = 42), ST485 (n = 25), non-typeable allelic profiles (n = 16) or ST651 (n = 14), with the remaining eleven STs represented by 3 or fewer genomes (see Microreact project). The Lac.2 operon was detected in nearly all GBS sequences of bovine origin (77/78), while only 13 % of human (17/130) or 5 % of food-derived (2/38) GBS sequences carried the locus. The *scpB-lmb* genes were not found in any of the bovine GBS genomes but were present in about 20% of the human and 10% of the food-associated genomes. Like *scpB-lmb*, six virulence genes associated with pilus structures (*srtC1*, *srtC2*, *srtC4)* and three pilin-related genes were associated with human or food isolates from Asia (see Microreact project).

Presence of *scpB-lmb* was never associated with carriage within our dataset (n = 0) (Table 1).

**Table 1:** Association between presence of scpB-lmb and health status of the human host.

|  |  | *scpB-lmb* |  |
| --- | --- | --- | --- |
|  |  | absent | present |
| Host health status | carriage | 20 | 0 |
|  | disease | 51 | 24 |
|  | missing values | 29 | 6 |

Consequently, the odds ratio and its upper confidence interval were infinite with a p-value of 0.003 (CI 2.1 – Inf), indicating that the odds of a GBS-CG103/314 genome carrying *scpB-lmb* increased strongly and significantly when isolated from a host with disease. This result remained consistent in sensitivity analyses when missing values for disease status were either coded as disease or carriage. Significance was slightly reduced in both sensitivity tests (p-values 0.007 and 0.005, respectively) and the effect strength reduced when missing values were coded as carriage (OR 3.8). Proportions of continents of origin among human derived genomes were in good agreement with overall proportion of continents in the full dataset.

#### Antimicrobial resistance gene patterns

ResFinder results at thresholds of 95% identity and coverage represented the overall findings from the three AMR databases well and were therefore used for subsequent analyses. Just over 30% of genomes had no AMR genes as detected by ResFinder. Across the remaining genomes, 15 AMR genes were detected, conferring resistance against multiple drugs (*optrA*), to aminoglycosides (*aph3..III*, *ant6.Ia*, *aadE*), macrolides, lincosamide (or the macrolides, lincosamides, and streptrogramin A and B (MLS) drug group; *lnuA*, *lnuB*, *mefA, ermB, msrD*), phenicols (*cat*, *catQ*) and tetracyclines (predominantly genes *tetM, tetO*, but also *tetL* and *tetS*). For details, please refer to the Microreact project. Nearly 50% of all genomes carried a single resistance gene, in most cases *tetM* or another tetracycline resistance gene, e.g., *tetO* (15% of genomes), especially in bovine isolates. This made tetracycline resistance the most common type of resistance across the dataset and DT analyses on the tetracycline resistance genes showed presence of tetracycline resistance genes from the time of emergence of the CGs (See Phylodynamic inference and Figure S8).

The prevalence of AMR genes differed between clades, host species and continents Network analysis and whole-genome-SNP phylogeny

Based on the network analysis, which retains the recombinant segments of genomes, ST103 showed a diffuse topology without a clear clustering pattern. By contrast, all but two ST314 genomes (40/42) clustered together (Figure 4) whereby three ST314 sub-clusters were identified. One, separated by a higher number of splits than the other two), consists mostly of human Asian sequences (see arrow in Figure 4). ST314 isolates appeared at the extremes of the network, confirming that the 7-gene MLST scheme does not fully represent the evolutionary history of genomes (81).

**Figure 4:**
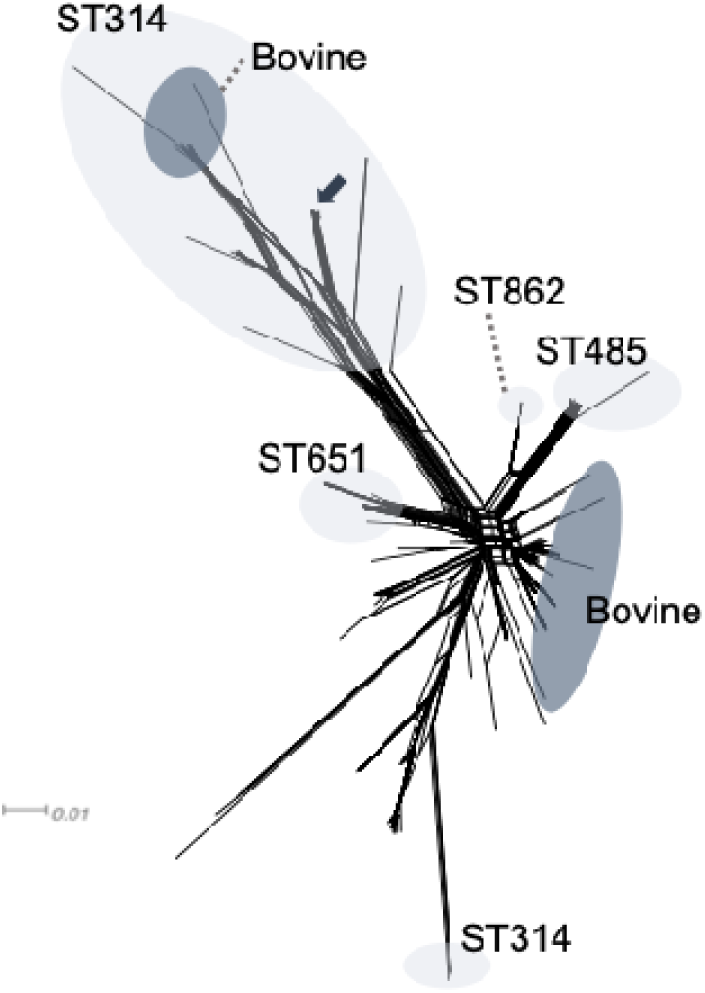
NeighbourNet network analysis of 248 Group B Streptococcus (GBS) genomes belonging to clonal groups (CG) 103 and 314 based on a whole core genome alignment including recombinant blocks. Ellipses indicate clades of predominantly (upper ST314 cluster and ST651 cluster) or exclusively one sequence type (ST) (light-grey) or genomes from the bovine host (dark-grey). Non-labelled clusters and sequences are predominantly ST103 of human or food origin. A small dark arrow points to a sub-cluster of ST314 sequences that originates more homogenously from human Asian sources than the remainder of the larger ST314 cluster it is nested within.

Closer to all other genomes but still in distinct clusters were ST485 and ST651, which were also relatively common in the dataset, and comprised of human and food isolates only, with human isolates dominating in ST485 and food isolates dominating in ST651 (see Microreact project). Genomes of bovine GBS clustered into two distinct parts of the network

The time-scaled recombination-filtered whole genome SNP-phylogeny was structured largely by ST, although ST103 encompassed several subclades, with three ST314 clades nested within it (Figure 5). These ST314 clades matched the distribution shown in the network analysis, i.e., one clade contained only human and food sequences from Asia and corresponds to the sub-cluster of the network graph marked by an arrow; one clade contained only two tips of ST314 matching the two sequences visible at the bottom of the network graph; and one ST314-enriched and bovine-dominated clade contained the remaining ST314 sequences of the network graph (Figure 4). This ST314-enriched clade showed a higher degree of topological uncertainty throughout analyses, with posterior probabilities supporting its placement below 70% for most runs. This is likely due to residual recombination in the alignment and corresponds to the network where ST314 genomes are separated by a higher number of splits (edge distance) from the remaining dataset (Figure 4). Food isolates clustered with human isolates, whereas animal isolates of non-bovine origin (pig, porcupine) clustered with bovine isolates. A demographic reconstruction showed no significant changes immediately following the year 2015, when the GBS outbreak in Singapore occurred (82), (Figure S6).

**Figure 5:**
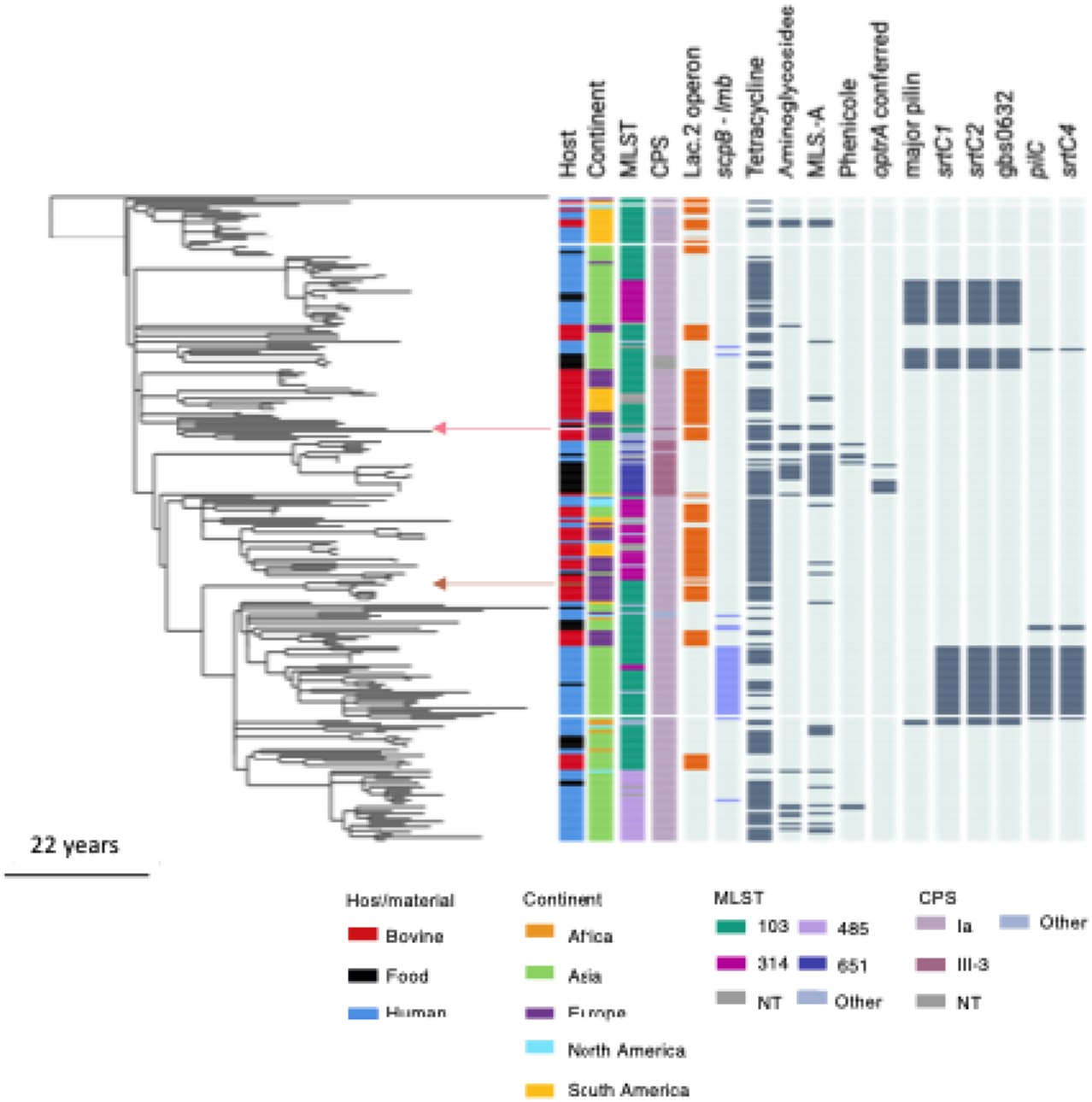
Time scaled maximum clade credibility tree based on Group B Streptococcus (GBS) genomes from clonal groups (CG) 103 and 314 (n = 245). Metadata are displayed as coloured panels on the right. Colours in the legend are given for host and continent of origin, as well as the largest groups of sequence types (ST) in the multi-locus-sequence-typing (MLST) system and capsular serotype (CPS). Minor STs and CPS, represented by fewer than 4 and 3 sequences, respectively, were grouped into “Other”. For a full list of associated metadata, see the Microreact project. Presence (colour) or absence (light grey) is indicated for the Lac.2 operon, scpB-lmb and antimicrobial resistance (AMR) genes, grouped by the drug class they confer resistance against (tetracyclines, aminoglycosides, the macrolide-lincosamide-streptagramin A (MLS-A) drug group, phenicoles and the gene optrA). The last six columns show data on virulence genes that showed differential presence/absence patterns across the dataset. The two sequences isolated from a pig (pink) and porcupine (chestnut) are highlighted with coloured arrows.

Across analyses with DT and MASCOT as well as different data subsets (Table S2), the time to the most recent common ancestor (TMRCA) was ∼80 years, placing the root of the tree at around 1943 (95% highest posterior density (HPD) interval [1927, 1958]), a time that also saw increasing diversification within major pre-existing human clades such as clonal complexes 1 and 23 (21). The genome-scale substitution rate for a genome length of 2,000,000 base pairs was 2.7x10^-07^ substitutions/site/year (95% HPD [2.3x10^-03^, 3.4x10^-07^]).

### Phylodynamic inference of host switching

Analyses using the structured coalescent approximation in MASCOT v3 inferred several host switches from human to bovine populations (Figure S9). Placement of host switches was very consistent between DT and MASCOT (compare Figure 7 further below and Figure S9) and both methods strongly favoured asymmetric transition models (analyses under symmetric priors largely failed to converge). MASCOT confirmed that food-derived genomes clustered with those from humans and showed no bifurcation as the food-association was present in single tips only (Figure S9). Population sizes for human-associated clades of CG103/314 were higher than bovine population sizes (Table 2). Host association of root and basal nodes of the tree was inconsistent throughout analyses.

**Table 2:** MASCOT-generated absolute migration rates (adjusted for backwards direction, left hand portion of table) and effective population size (N_e_, right hand portion of table) for Group B *Streptococcus* (GBS) clonal group 103/314. Results are shown for GBS populations associated with human or bovine hosts under the assumption of constant population size. HPD - highest posterior density (interval).

| | Average host switches/year | 95% HPD | Host | $N_e$ | 95% HPD |
| --- | --- | --- | --- | --- | --- |
| **Human to Bovine** | $1.9 \times 10^{-2}$ | [5.8E-3, 3.3E-2] | Human | 672 | [490, 876] |
| **Bovine to Human** | $5.3 \times 10^{-4}$ | [1.8E-6, 1.5E-3] | Bovine | 280 | [161, 406] |

### Geographical origin and spread of clonal groups 103/314

When stratifying data by continent-host combination, major monophyletic clades included multiple hosts and continents but smaller clades were mostly limited to a single continent-host combination. Transitions between continents seem to have happened predominantly prior to the 2000s (Figure 6). DT analysis of migration rates using selection based on Bayesian stochastic search variables suggests migration of European bovine isolates into humans and cattle across all continents (Figure 6 and S10), including multiple introductions into South America. One of the South American sub-clades leading to multiple tips transitioned earlier than the others (yellow arrow) and both DT and MASCOT phylogenies inferred bovine association preceding this node. There is one even earlier introduction represented by a single tip and given the early inferred timepoint, a lack of available samples might be obscuring the subsequent evolution of this clade. The acquisition of *scpB – lmb* appears to have happened after the transition into Asia (violet arrow, *scpB-lmb* clade shaded box). A continent level DT phylogeny (not focusing on host association, Figure S11) tentatively suggests that most transitions into Africa happened from Asian sources.

**Figure 6:**
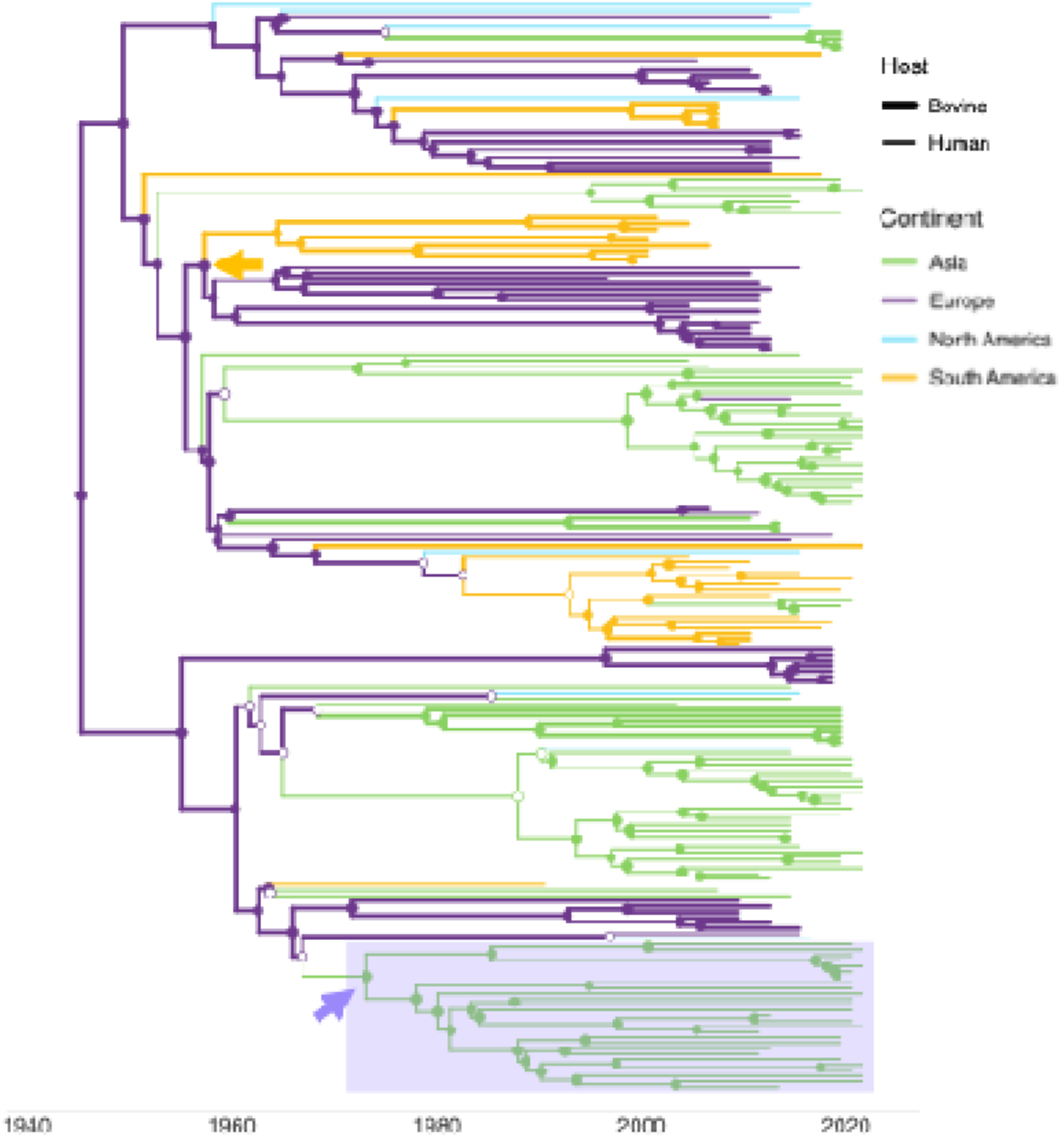
BEAST-1 discrete phylogeographic analysis of subpopulations of Group B Streptococcus (GBS) clonal group 103/314 genomes (n = 199) stratified by host-continent combination. Genomes from Africa are excluded because of an insufficient number of sequences. Time scaled phylogeny with line thickness according to host species (shown in legend) and branches coloured by continent. Node colour fill indicates a support of posterior probability > 0.9 for association with this host/continent combination (empty nodes indicated node support below 0.9). Arrows indicate an early transition into South America (yellow) and acquisition of scpB-lmb after transition into Asia (violet).

### Marker genes and phylodynamic inference of ancestral presence/absence

The bovine marker Lac.2 was inferred as being present from the emergence of both CGs, despite the association of basal nodes with human hosts in this analysis, a discrepancy that could potentially be due to dataset imbalance. The Lac.2 operon was subsequently lost in several human-associated GBS clades before the 1960s and either retained or regained in all bovine clades, while two human clades also retained the Lac.2 operon until after the 1960s (Figure 7 orange boxes). One of these clades contains predominantly human GBS genomes from Brazil, the other has no predominant metadata trait.

**Figure 7:**
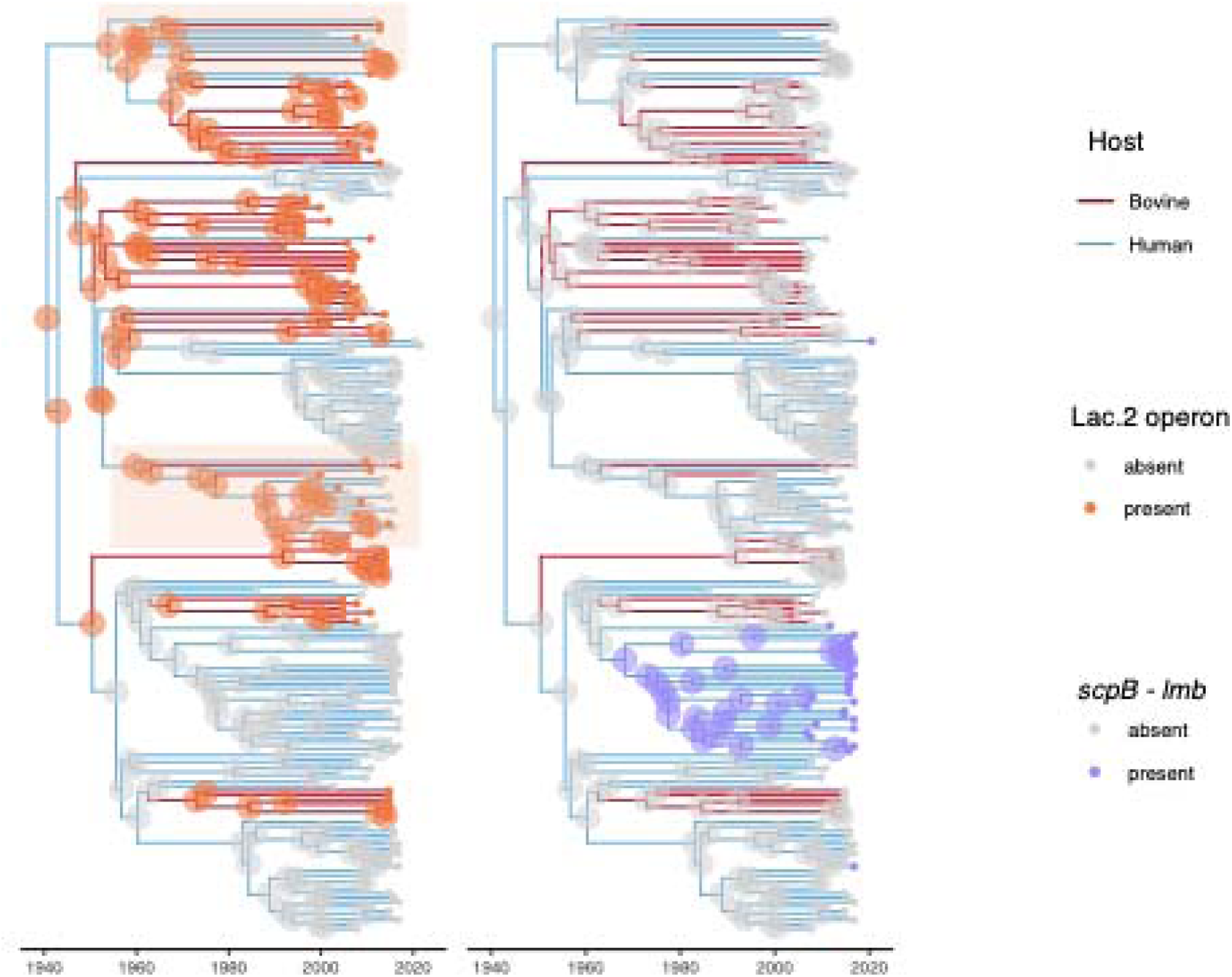
Time-scaled maximum clade credibility tree based on the Group B Streptococcus (GBS) dataset of clonal groups 103 and 314 visualising host association and gene loci. The trees were constructed through discrete-trait mapping of host marker genes and host of origin in BEAST-1. Data were obtained from a dataset with available metadata on year of isolation but excluding genomes from pigs, porcupines and food (n = 205 genomes included). Branches are coloured by inferred host association; nodes are coloured by inferred presence (coloured) or absence (grey) of genes, while tips are coloured according to known presence or absence of the same genes: left – Lac. 2 operon as marker of bovine host association and right – scpB-lmb as marker of human host association (19); the clades coloured in orange retained the Lac.2 operon in human associated lineages beyond 1960.

The *scpB-lmb* genes, which are markers of human host adaptation, were mostly absent from the tree and only acquired on a few occasions; two of these are lineages represented by a single tip (Figure 7). A single clade acquired the *scpB-lmb* transposon around the 1970s. This clade comprises exclusively of human GBS genomes from Asia, mostly of ST103. Results from the test of association between presence of *scpB-lmb* and disease status of the human host did not present visually clear within the phylogeny: genomes with available metadata reporting (any) disease were distributed throughout the tree without a visually obvious correlation with *scpB-lmb* presence or absence (see Microreact Project).

## Discussion

We compiled a dataset from multiple continents of two multi-host CGs of GBS, 103 and 314, which have been found primarily in humans and cattle, with rare reports from other host species. We characterised their population structure, assessed AMR, virulence and host-associated gene content, and investigated whether phylodynamic analyses can reveal trends in transition directions between host populations. As a result of diversity in surveillance or research efforts, our genomic dataset included bias in terms of geographic origin, host species, and time of collection of GBS isolates. In particular, Asian CG103 of bovine origin is underrepresented in the genomic dataset compared to the literature (27, 33, 83), whilst the peak in bovine GBS isolates from Europe represents a concerted research effort in Northern Europe around 2010 (20, 25, 26). The lack of human genomes from Europe appears to represent the true, low prevalence of CG103/314, at least in the Netherlands and Denmark. In both countries, ST103 was detected only once among collections of thousands or hundreds of human isolates, respectively, covering maternal carriage, neonatal infectious disease, and adult GBS disease across multiple decades (personal communications M. Bijlsma, Amsterdam Medical Centre, Amsterdam, the Netherlands, and U.B.S. Sørensen, Aarhus University, Aarhus, Denmark, 2025). The 2015 GBS outbreak in Singapore was the motivation for comprehensive sampling efforts and the origin of many of our Singaporean GBS genomes, including those from food (82, 84, 85). Otherwise, it is not known whether the apparent emergence of GBS CG103/314 in humans in Asia represents sampling effort or the evolution and epidemiology of the clade.

Imbalance in sampling effort exemplifies a common obstacle in the study of One Health pathogens, that is, bias towards the research of pathogens and strains that affect human health or economically relevant animal species (86, 87). The true prevalence across host species of a given multi-host pathogen or lineage is often obscured by a lack of surveillance effort, e.g., in wildlife or in healthy individuals that carry the organism. GBS, for example, is primarily studied in neonates and pregnant women, leaving the prevalence and population dynamics in men and non-pregnant women poorly characterised. Additionally, sampling, surveillance and pathogen characterisation, especially the use of costly tools such as whole genome sequencing, extends the bias towards immediate threats as well as higher income countries (88–90). As an example, ST248 was very rare in our genomic analysis (n = 1), whereas ST248 was more common than ST314 or ST103 among GBS isolates from cattle across different regions in Colombia (23, 91). Attempts to generate more genomic data from the Colombian ST284 isolates were affected by logistic hurdles such as freezer breakdown and loss of isolates, lack of staff or funds, and reallocation of resources during the Covid-19 pandemic.

### Prevalence of AMR differs by host and region

In this global CG103/314 dataset, a variety of AMR profiles were observed across continents and host species. Asian GBS genomes, which were primarily of human origin, exhibited the highest frequency and diversity of AMR genes. In human isolates from Europe, the prevalence of AMR genes was lower than in human isolates from Asia, but it was higher than in bovine isolates from Europe. Conversely, the prevalence of AMR genes in human isolates from South America was lower than in bovine isolates from that continent. Although we did not measure associations with antimicrobial use, those patterns may reflect the existence and enforcement of antimicrobial use policies in different regions, and the prevalence of GBS in different host species. For example, antimicrobial use in China is high in comparison with Europe, which contributes to differences in AMR emergence (92). Likewise, antimicrobial use in animal production in Thailand is high, with demonstrated impact on prevalence of AMR (93–96). The detection of lower levels of resistance (here: number and diversity of resistance genes) in human than in bovine GBS genomes from Brazil aligns with findings by Pinto et al. (2013), who reported ∼ 20 to 30% resistance to clindamycin and erythromycin in bovine isolates, compared to ≤ 4% in human isolates (97). Despite attempts to limit antimicrobial use in livestock through regulation and enforcement, use of broad-spectrum antimicrobials without proper technical advice is still quite common in segments of the Brazilian dairy industry, which may contribute to this difference (98–100). Tetracycline resistance gene presence in human GBS is so common that it was once thought to be a marker of human host adaptation (21), an association that has since be dispelled based on genome wide association studies across host species (19).The presence of *tetO*, in particular, occurs more commonly in bovine GBS than in human GBS in the current study, as previously observed for temporally and spatially matched human and bovine isolates from the USA (101). Group B *Streptococcus* is inherently resistant to kanamycin (low level) but presence of aminoglyside resistance genes is rare in GBS (102, 103). Here, *aad,* a gene encoding an aminoglycoside nucleotidyltransferase (also referred to as ANT(6)-Ia) was detected in just under 10% of genomes of the current dataset, most of them from Asia.

The highest diversity of AMR genes was observed in the ST651 clade, encompassing ST651, ST862, and ST1983 from humans and food (mostly products from pig) sources, which mostly originated from Asia. A study in Hong Kong (39) found ST651 and ST862 to be the predominant GBS types among pig-derived isolates, particularly from pig tongues. The detected AMR profiles and an absence of pilus genes from GBS ST651/862 in their study is consistent with our findings. Results from Hong Kong, and the fact that we found six AMR genes in a pig isolate from Europe, suggest that further research into the prevalence and role of GBS and AMR in pigs may be needed.

### Population structure of clonal groups 103 and 314

Both the network graph and whole-genome SNP-phylogeny showed clustering by ST, although polyphyletic. We observed a separation of the ST314 clades in both the network graph and the SNP-based phylogeny, whereby the network approach specifically accounts for recombination. The phylogeny separated the genomes into three distinct clades which broadly mirrored the structure of the network graph. However, the phylogeny separated the ST314 sub-clades far from each other by placing them each into one of the two major monophyletic clades and even nesting one of them within a larger ST103 context. In contrast, in the network graph most of the ST314 genomes cluster more closely together, aligning better with expectations for ST groupings and raising doubts about the appropriateness of a SNP-based phylogeny to reflect the evolutionary history of these CGs accurately. Studies in other species of bacteria indicate that intra- and intergenic recombination can affect housekeeping genes, such as those used in the MLST scheme (104, 105). Evidence for this in GBS has been previously described (81) including in housekeeping genes. However, it should be noted that in the context of the global GBS population, CG103/314 has a low rate of recombination, in particular when compared with other host generalist clades (19). We strongly support including recombination information into analyses on GBS population structure regardless of CG to allow the consideration of any horizontally inherited genomic features. For several genomes, an ST could not be assigned because no allele number was assigned to the *glcK* locus. This has been observed in GBS analyses previously and is likely connected to an indel event resulting in truncation of the gene (58, 106). While raising doubt about the role of *glcK* as housekeeping gene and affecting ST nomenclature, core or whole genome analysis still enables placement of those genomes in the networks or tree graphs.

### Phylodynamic analyses

Because bovine GBS genomes from Europe and human GBS genomes from Asia dominate the dataset, the outcome of phylodynamic analysis of CG103/314’s early host association, population size, migration direction and rates may be affected by bias. In attempts to mitigate these biases, we conducted a series of sensitivity analyses using two phylodynamic methods (DT and MASCOT), different data subsets, subsampling to balance out traits (supplementary material Section 2) and varying trait-groupings such as continents vs. global regions. Despite these efforts, results for early host association remained inconclusive. Similar problems have been described for attempts to determine the origins of ST283, a multi-host strain of GBS that primarily affects humans and fish and has transitioned between the two groups of hosts on multiple occasions (85). In our MASCOT analyses, CG103/314 primarily transitioned from humans to bovines and primarily before the 2010s, aligning with larger human-associated GBS population size estimates. These findings extend earlier research by Richards et al. (2019), who described that human-to-bovine transitions are more common than bovine-to-human transitions in GBS evolution, but who had limited access to CG103/314 genomes (107).

Historical data are available around potential transitions between continents but do not necessarily provide clarity either. On the one hand, exports of Dutch dairy cattle to Brazil and other Latin American countries are documented, supporting the possibility of European-to-South American transmission (108). On the other hand, cattle and agricultural products were imported from South America, notably Argentina, to Europe after World War II, which devasted European economies (109). Despite large scale imports from South America to Europe, we do not see this direction of transition in our phylogeographic analysis, which may support a European origin of CG103/314. However, sensitivity analyses using subsampling to balance geographical and temporal bias indicated an origin of CG103/314 within the Americas and the first known CG103/314 genome originates from a human in Brazil in the 1990s (28, 29, 58). As for ST283, more geographically balanced sampling and regard to demographic changes over time - where known - would allow for more confident conclusions regarding the origins of CG103/314 (110, 111).

### CG103/314 carries markers of bovine rather than human host-adaptation

GBS host adaptation to bovines and humans has previously been found to be strongly linked to presence of the Lac.2 operon and the *scpB–lmb* gene pair, respectively (5, 19). In our dataset, the Lac.2 operon was present in all bovine derived CG103/314 genomes, in agreement with its role as marker and functional facilitator of bovine host association, but it was not restricted to bovine GBS genomes. This locus, which comprises 9 to 11 genes, is located on a putative integrative conjugative element, suggesting acquisition via horizontal gene transfer (HGT), and plays a key role in lactose fermentation, facilitating nutrient utilisation in the bovine mammary gland (5, 112). Our phylodynamic analyses inferred presence of the Lac.2 operon early in the evolutionary history of CG103/314. However, as the phylogeny was built from vertically inherited SNPs and the presence/absence of loci was mapped as a discrete trait onto this fixed topology, such inference should be interpreted cautiously for elements acquired through HGT. In DT tools, trait changes are modelled in a similar manner to point mutations (113) but that approach does not account for HGT. This suggests the need for other tools than the BEAST framework for modelling its gain and loss onto a phylogeny (114, 115).

*ScpB-lmb*, a set of virulence factors co-located on a single transposon, is crucial for invasive disease in humans (116–118). Existing literature on the broader GBS population suggests a strong association between the *scpB-lmb* gene pair and human hosts (19, 119). In line with this association, *scpB-lmb* was absent from bovine GBS genomes. All sequences containing *scpB-lmb* originated from cases of human clinical disease rather than asymptomatic carriage. However, the gene pair was absent from most human GBS genomes in CG103/314, including from all carriage isolates and a proportion of clinical isolates. This absence may, at least in part, explain the low virulence of CG103/314 in humans (28, 120). The exception to this is ST485, which has been linked to severe disease in Asia (30), although the ST485 clade was not associated with the presence of *scpB-lmb* in our dataset.

### Emergence in new host species illustrates ongoing evolution

In food isolates, two potential patterns of transmission were observed. On the one hand, the distribution of GBS genomes from food isolates within human clades suggests that GBS detection in food may result from human contamination during handling, as previously described for other GBS sequence types (121, 122). On the other hand, there is a sub-clade of ST651 that does not include human isolates but consists of eleven food isolates, originating from aquatic species (tilapia, threadfin and squid; n = 7), poultry (n = 1), and pigs (n = 3).

Poultry are not a known host of GBS, nor are any other bird species, and detection in a chicken gizzard could be due to ingestion of food of other animal origin, or results from contamination during handling. Members of CG103/314 have been reported from diseased fish (ST103, Brazil) and prepared fish purchased at market (ST651, Hong Kong) but such reports are very rare (123, 124). Based on the available metadata (year of isolation), tilapia were sampled in a single year but it was not possible to determine whether they represented a single batch of fish from a single farm or market stall. Difficulties in determining the origin of GBS in seafood have been reported before and are compounded by accumulation along the processing chain from catch to market and retail (16). The only animal species to yield ST651 on multiple independent occasions, as evidenced by year of isolation, was pigs. Combined with the fact that ST651 was the predominant type in pig offal in Hong Kong, this suggests that pigs may be an emerging host for GBS. Emergence of a GBS lineage in a new host species has recently also been described for elephants (125) and porcupines (see below and Figure S9). If GBS ST651 causes oropharyngeal colonisation in live pigs, as suggested by data from Asia, (39, 126), this may result in introduction into the food chain via healthy animals, particularly from tongues and tonsils. Reports of GBS in live pigs in other parts of the world are scarce, with unclear links to disease (127, 128). Our genomic analysis included a single isolate from a pig that died with pneumonia due to GBS, probably acquired through consumption of raw milk whey (38).Whether or not a pig with clinical disease would enter the food chain depends on local approaches to food safety.

Notably, we were able to include the genome from an early instance of GBS causing respiratory infection in a wild porcupine in Italy (38). Garbarino et al. proposed the existence of a distinct porcupine-specific ST103 lineage. Within the current dataset, the porcupine genome appears adjacent to, but on a separate branch from, the clade of Italian bovine genomes in both the core-genome SNP phylogeny and the network graph. Although a single isolate cannot differentiate between incidental spill-over and establishment of a reservoir in a new host, its placement supports the possibility of the latter. For porcupines, consumption of contaminated bovine milk seems an unlikely route of exposure. Instead, presence of CG103/314 in faecal or environmental samples on dairy farms (22, 24) may facilitate its spread to a wider range of hosts, including wildlife. Bovine-to-wildlife spread via faeces or the environment has previously been described for *Mycobacterium bovis* in the UK. It is currently deemed rare but may have seeded *M. bovis* infections in badger populations at a time when fewer control measures were in place for bovine tuberculosis (129). A similar phenomenon could theoretically lead to emergence of GBS in wild porcupines.

## Conclusion

The current study offers a comprehensive characterisation of the majority of available CG103/314 genomes, two intriguing CGs within GBS. Leveraging a multi-continent dataset, we were able to show its broad distribution and some unexpected findings, such as the absence of human virulence markers in the majority of human-derived strains, as well as AMR and virulence patterns that vary by ST, country level and host species. Detections in symptomatic pigs and porcupines suggest these CGs may have the capacity to infect a broader range of hosts than currently recognised. Profiles of higher and more diverse AMR and virulence gene presence were observed in distinct clades, predominantly from Asia.

Together with repeated reports of CG103/314 causing human disease in China within the last decade, questions arise about possible evolution of new sub-lineages under selective pressure. The example of these CGs highlights the relevance of progressing past the anthropocentric perspective and taking a One Health and collaborative cross-border approach during the genomic monitoring of multi-host zoonotic pathogens.

## Acknowledgements

We thank the organisers of the third International Symposium on *Streptococcus agalactiae* Disease (ISSAD; Brazil, 2023) and the Microbiology Society for inviting us to present this work. We thank the IT department of the University of Glasgow for their assistance in data handling. We thank Dr. Lucy Weinert of the Cambridge Veterinary School and Dr. Paul Johnson from the University of Glasgow, as well as the members of the Epidemiology, Economics and Risk Assessment (EERA) group, especially Dr. Bryan Wee and Dr. Jamie Gorzynski, at the Roslin Institute, University of Edinburgh, for their valuable advice and input on this work. We thank Marco von Zwetselaar for his support in the implementation of assembly pipelines.

## Ethical approval

Ethical approval for processing of Singaporean samples was granted by The National Healthcare Group Domain Specific Review Board (Ref DSRB 2018/01010).

## Contributions

A.H., R.N.Z., C.C., T.F., R.B. and S.L. designed research; A.H. performed research; T.B., S.C. M.R., C.C-A. W.S., S. A-N., N.N.P., T.P., L.O., contributed isolates or genomic data; A.H. analysed data; and A.H., R.N.Z., C.C., T.F., R.B., S.L. and T.L. wrote the paper. All authors read and approved the final version of the manuscript.

## Microreact Project

URL: https://microreact.org/project/qdpMsBtAX8FJmTXVZm85AZ-gbscg103314hilbigetal Link: MicroreactProject-gbscg103314hilbigetal

