## Supplementary Materials for "Home is where the host is: Evolutionary history of geographic spread, host switching, and adaptive genomic signatures in two generalist Group B *Streptococcus* clonal groups"

##### 1. Detection of host marker genes

Detection of marker genes was carried out using custom libraries (1). The *scpB-lmb* library contained six variants of *scpB* and two variants of *lmb*. While some genes are shared between the lac.1 and Lac.2 operon, *lacE* and *lacG* are unique to the Lac.2 operon and thus were used for detection (2). The *lacE*-*lacG* library contained six variants of *lacE* and eight variants of *lacG*. Presence of a locus was called if there was at least one variant of each gene with > 90 % of both identity and coverage among all contigs of an assembled genome.

##### 2. Overview: Subsampling approach to mitigate dataset bias

As an additional attempt to mitigate the effects of sampling bias, we employed a subsampling approach. Ideally, aside from host species and time-interval, clonal group and continent of origin would also have been considered as traits to subsample by. However, it was not possible to populate categories stratified by all these traits with a number of genomes sufficient for meaningful analysis. We thus chose to conduct two separate subsampling analyses, one for the host association and one for host-continent association. Because of the under-representation of human associated genomes during the early sampling years and over-representation in later years, both subsampling schemes used a dataset where the time range was limited to 2004 and 2019, during which both human and bovine GBS genomes were available. To maximise the usable data, food-associated genomes were coded as human. All food-associated genomes had clustered with and evenly throughout human subclades (see section 2, Figure S9).

###### 2.1. Methods: Subsampling for ancestral state reconstruction of host association

One clade of predominantly CG 103 was chosen from the phylogeny for host-subsampling (n _genomes_ = 76). This clade inherently displayed an adequate degree of balance for host, continent and year-range. The years of 2004 – 2019 were sectioned into 3-year intervals and for each interval a maximum of 3 genomes per host was sampled at random. If fewer than 3 genomes were available, all available genomes were sampled. This process was repeated 10 times to produce 10 subsampled alignments of n_genomes_ = 27 genomes.

###### 2.2. Methods: Subsampling for ancestral state reconstruction of host and continent association

The entire dataset, truncated to the years 2004 – 2019 and excluding African genomes (n_genomes_ = 163), was used for host/continent-stratified subsampling. The main difficulties in non-subsampled attempts to run host/continent stratified analyses had been the use of too many trait populations (demes) and the imbalance in genome number within these demes. To mitigate this effect, we pooled European and Asian genomes to represent Eurasia, and North- and South American genomes to represent the Americas. The years 2004 – 2019 were sectioned into 5-year intervals and for each interval a maximum of 3 genomes per combination of host and global region was randomly sampled. If fewer than 3 genomes were available, all available genomes were sampled. This process was repeated 10 times to produce 10 subsampled alignments of n_genomes_ = 40.

Both sets of subsampled alignments were run in MASCOT v.3 (3). To counteract the reduction in sequence data information, priors for substitution rate and population size were adjusted to be more informative. Parameters across all replicates were summarised in R (R-Core-Team, 2023), 95% HPD intervals were calculated using the R package ‘coda’ (4).

###### 2.3. Results: Subsampling for ancestral state reconstruction of host association

Within the context of the maximum clade credibility tree produced by MASCOT using the full dataset, the root of the subsampled clade was associated with human hosts with a high posterior probability of > 98 %. However, all ten analyses using time and host balanced subsets of this clade resulted in the root and early ancestral nodes being associated with bovine hosts, at posterior probabilities (for both root and early nodes) between 81 – 99 % posterior probability (mean 93 %, median 97%).

###### 2.4 Subsampling for ancestral state reconstruction of host and continent association

There was no consensus about the host associated with root and early nodes. More surprisingly, as it stands at odd with the other results, the global region associated with the early regions of the tree was consistently the Americas. Out of 10 subsets 9 converged and out of these 9 analyses, 4 showed human-associated GBS genomes from the Americas at the root, 4 showed bovine-associated GBS genomes from the Americas and one run inferred almost equal probabilities for either.

### Supplementary Figures

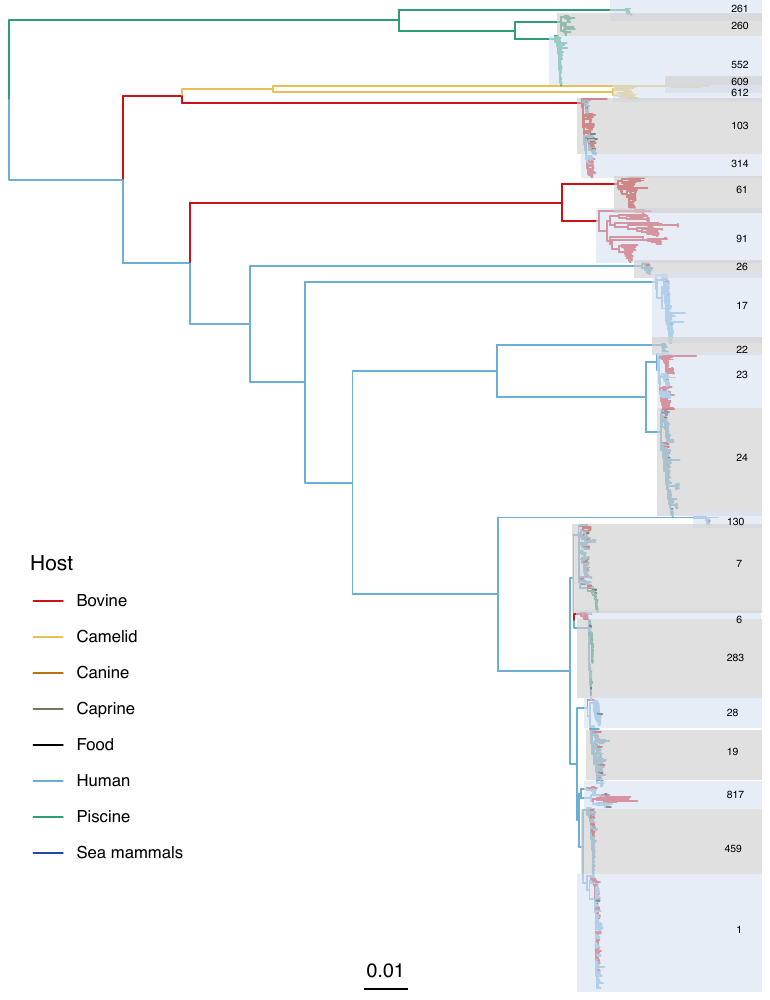

**Fig. S1:** **Maximum-likelihood tree based on a global dataset of the Group B Streptococcus (GBS) population visualising ancestral host association.** Figure adapted from Crestani et al. 2024, please see publication for details. In brief, the tree was constructed using IQ-Tree and discrete-trait mapping of host association was carried out using the ACE function of R-package ‘Ape’ but was not published along with the other results. The dataset (n = 1254) is described in detail in the cited publication. Clades are annotated with their corresponding clonal group; alternating colours of boxes around clades (grey/blue) were used to improve visual distinction between boxes, colours bear no meaning.

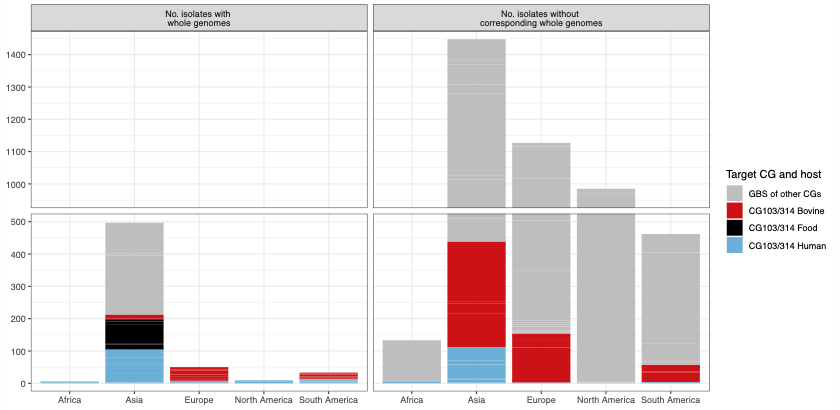

*Fig. S2: Collections of Group B Streptococcus isolates that contained at least one isolate of clonal groups 103 or 314. Depiction is separated into those isolates that were sequenced (left) and those that, according to a literature research conducted, remain without whole genome sequencing (right). Isolates belonging to either of the two target CGs, sampled from human, bovine or food sources are shown in colour, whereas all other CGs are represented in grey. An axis break is introduced between 500 and 1000 isolates for better legibility.*

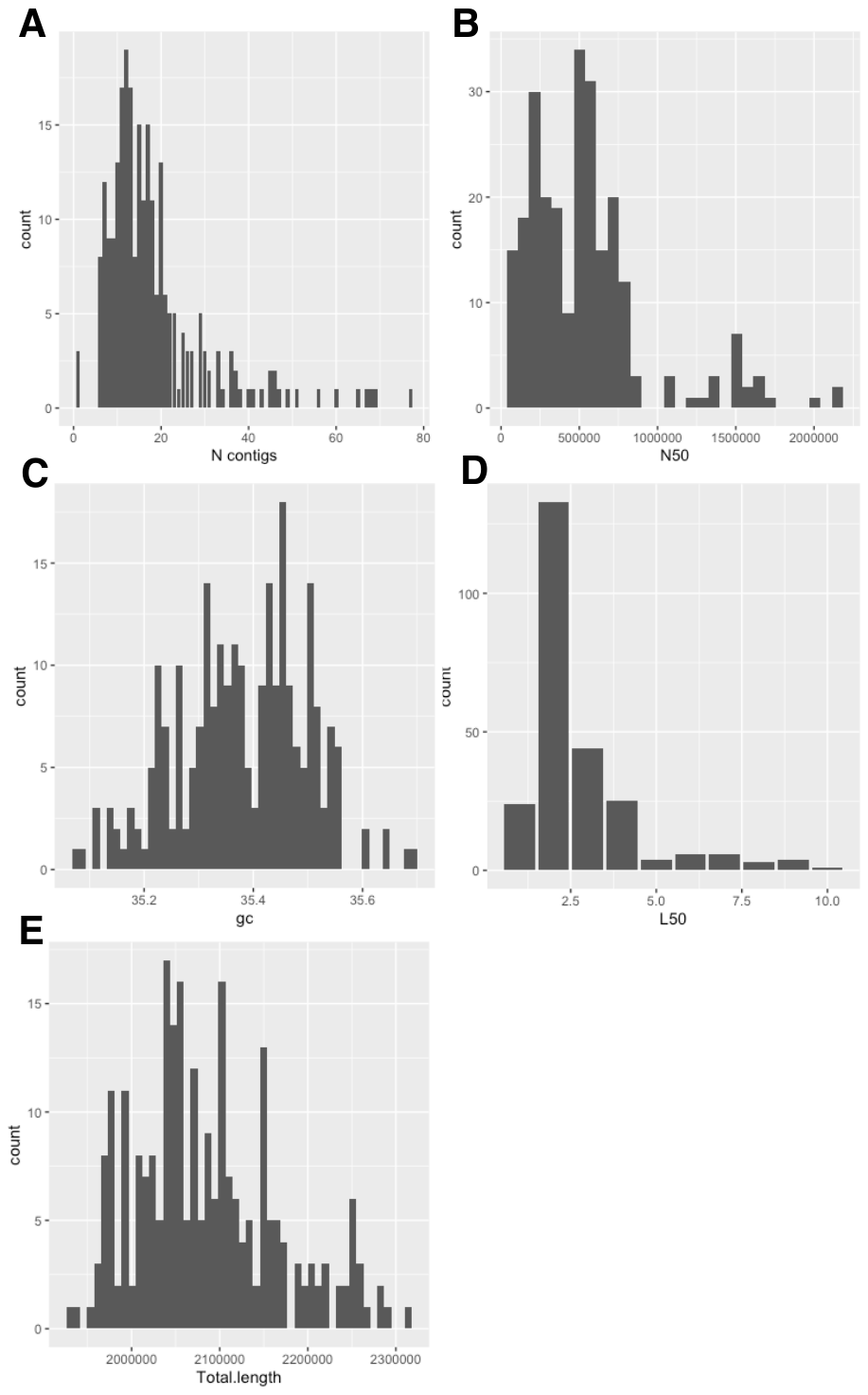

**Fig. S3:** **Distributions for metrics for quality assessment of assembled Group B Streptococcus (GBS) genomes (n = 250).** A through E: Number of contigs, N50, Guanine-Cytosine (GC) content, L50, and total length of assembled genome in base pairs.

**Fig. S4:** **Maximum likelihood phylogeny based on the alignment of all genomes of Group B Streptococcus (GBS) clonal groups (CG) 103/314 (n = 248).** The tree was generated with IQTree and recombination detection was done using Gubbins. Coloured panels to the right show continent and host of origin as well as sequence type (ST) based on multi-locus sequence typing (MLST). Block indicates recombination. Red signifies inferred ancestral status (occurred at a non-terminal node) and blue if they only affect one isolate/the tip. Boxes indicate the ST651/862 (middle box) and the two ST314 clades (top and bottom boxes).

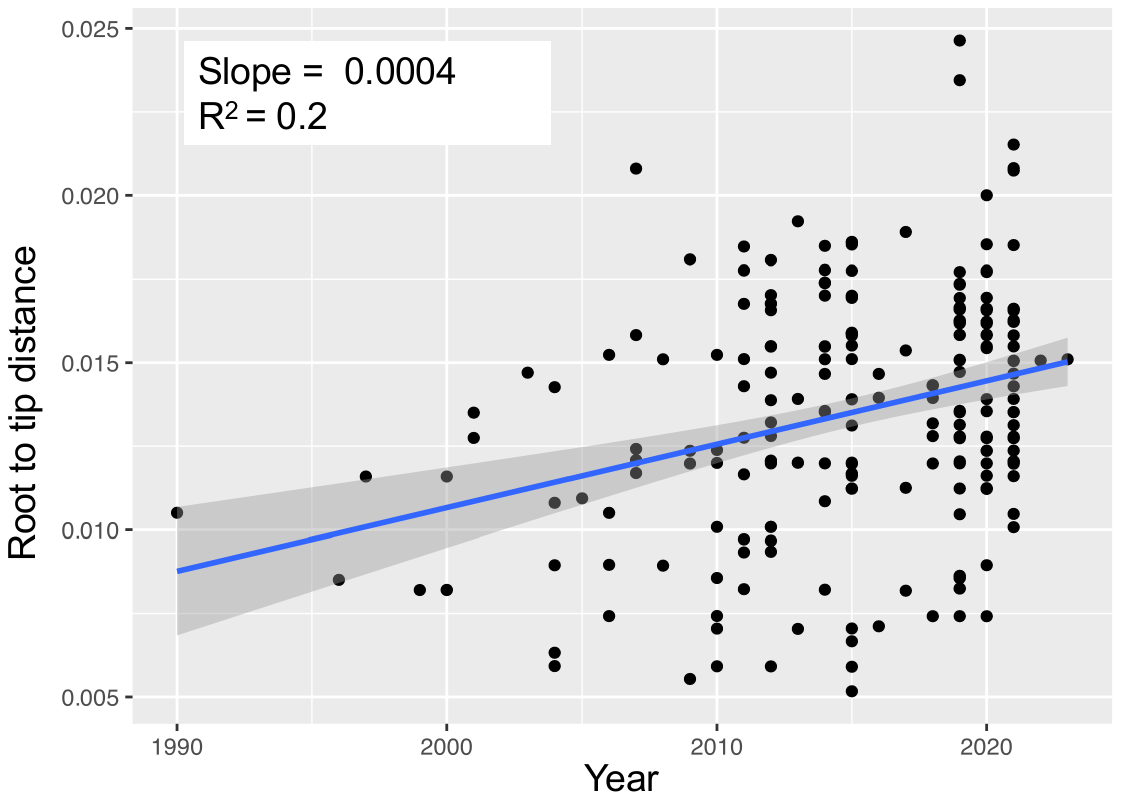

**Fig. S5: Root-to-tip linear regression analysis generated from a neighbour-joining-tree that was based on the alignment of all genomes of Group B Streptococcus clonal groups 103/314 for which year data were available (n = 245)**. One phylogenetic outlier has been removed. P-value < 0.05 for the slope coefficient. Blue line indicates the regression line, the grey shading indicates the confidence interval. Root to tip distance on the y-axis is measured in nucleotide substitutions.

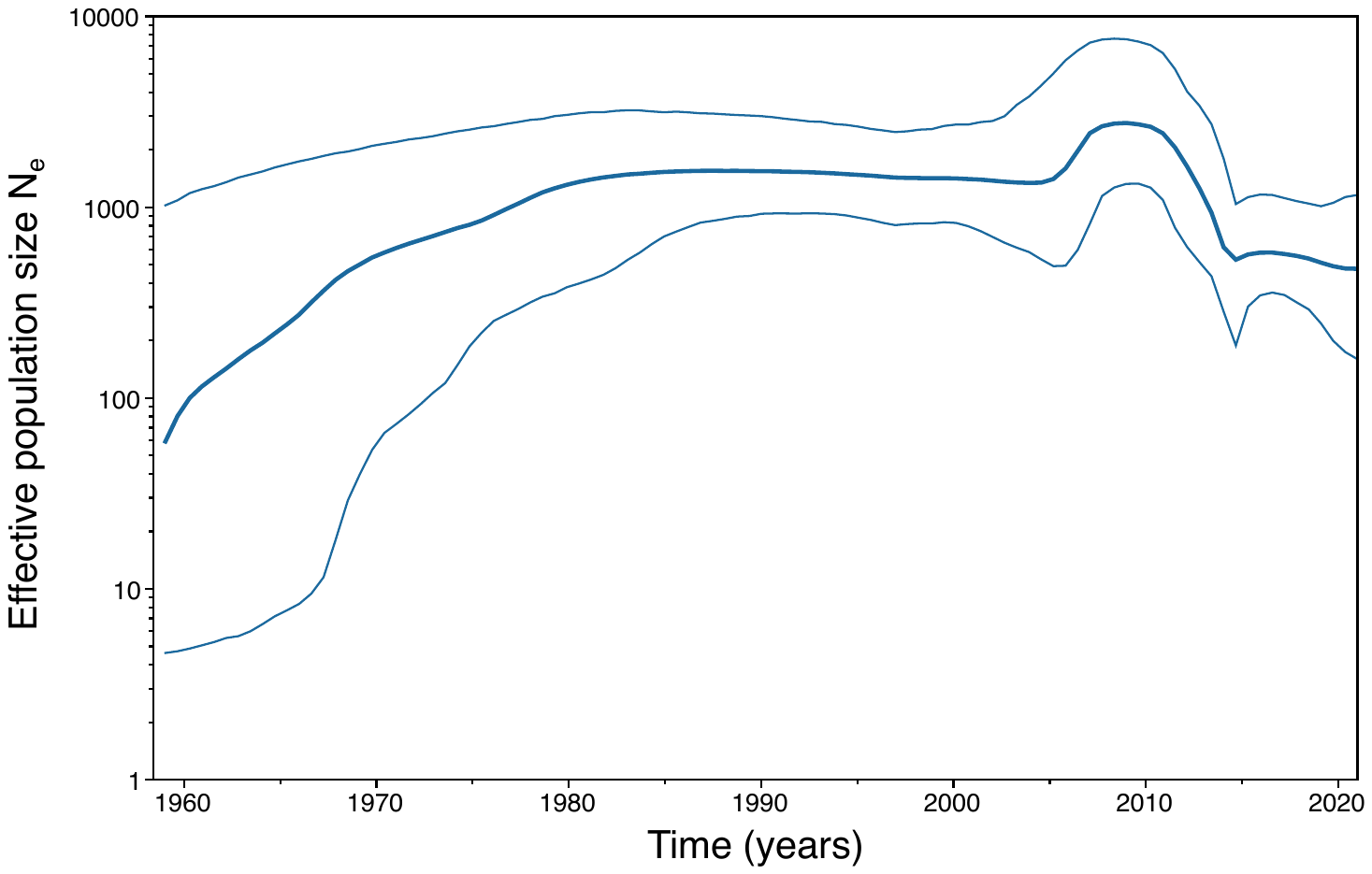

**Fig. S6:** **Bayesian Skyline reconstruction of demographic history generated in Bayesian Evolutionary Analysis Sampling Trees (BEAST)**. This is based on the clonal groups (CG) 103/314 dataset including all genomes with available information on sampling year (n = 245). Time is plotted on the x-axis against effective population size (N_e_) on the y-axis.

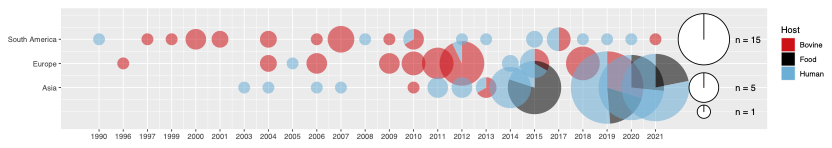

***Fig. S7: Visual representation of geographic and temporal coverage of the CG103/314 whole genomes in the dataset****. Pie sizes representing the number of CG103/314 genomes with pie proportions and colours indicating the host or material of origin. Genomes from Africa and North America were excluded from the visualisation due to insufficient data.*

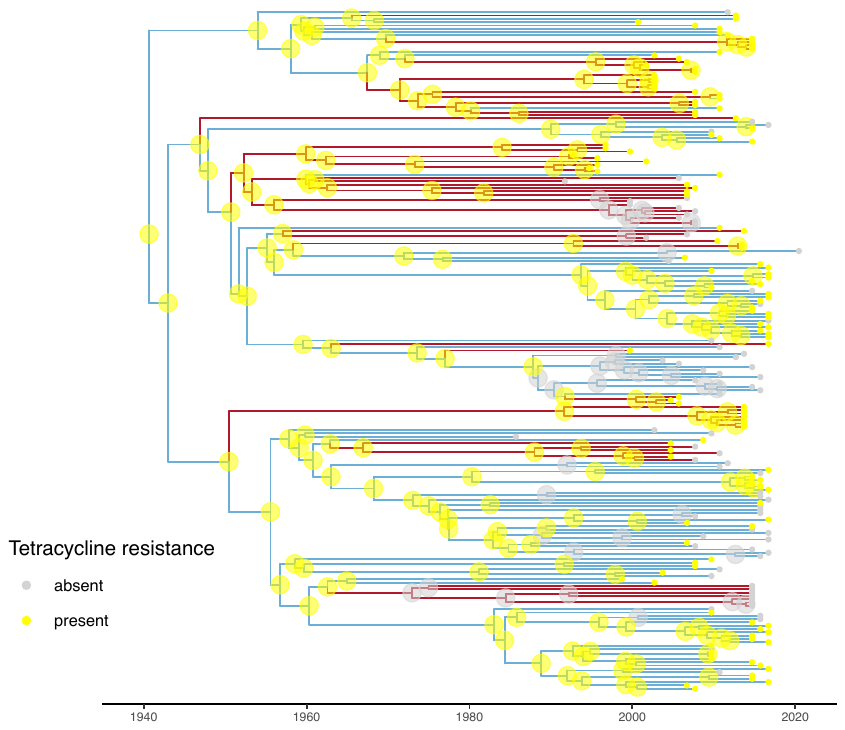

**Fig. S8: Time-scaled maximum clade credibility tree based on the alignment of all genomes of Group B Streptococcus clonal groups 103/314 dataset, for which year data were available (n = 245).** The phylogeny was generated using discrete trait mapping in Bayesian Evolutionary Analysis Sampling Trees (BEAST)-1. The branches are coloured by inferred associated host (bovine red, human blue). The presences of (any) tetracycline resistance gene (in this dataset we detected either tetM, tetO, tetS or tetL) is represented as a small solid tip circle and the inferred presence as a large transparent node circle, yellow for present, grey for absent.

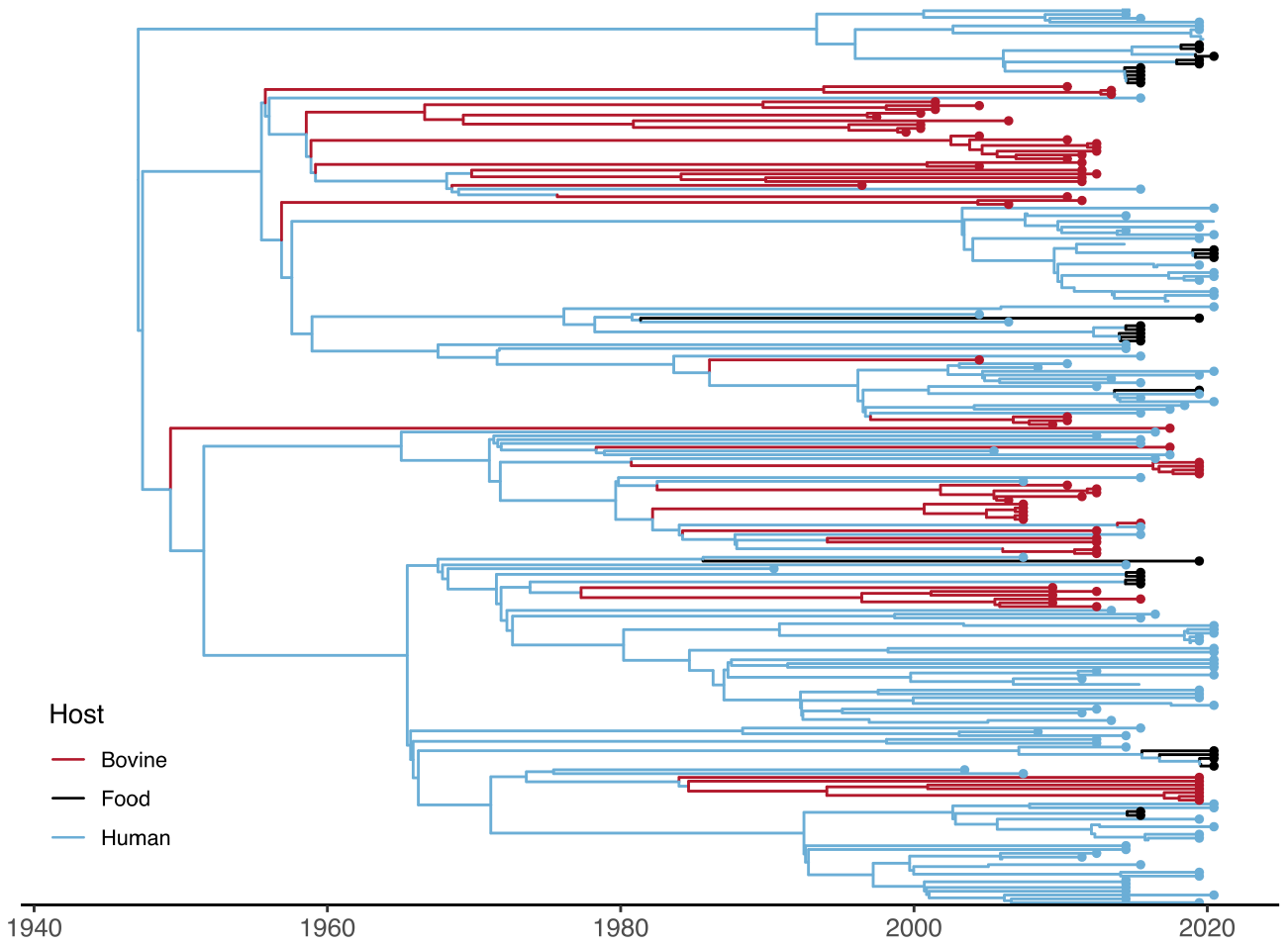

**Fig. S9:** **Time-scaled maximum clade credibility tree based on the alignment of all genomes of Group B Streptococcus clonal groups 103/314 for which year data were available and excluding pig and porcupine genomes (n = 243).** The tree was generated with the Marginal Approximation of the Structured Coalescent (MASCOT) v.03 as implemented in Bayesian Evolutionary Analysis Sampling Trees (BEAST) – 2. The branches are coloured by inferred ancestral-host association.

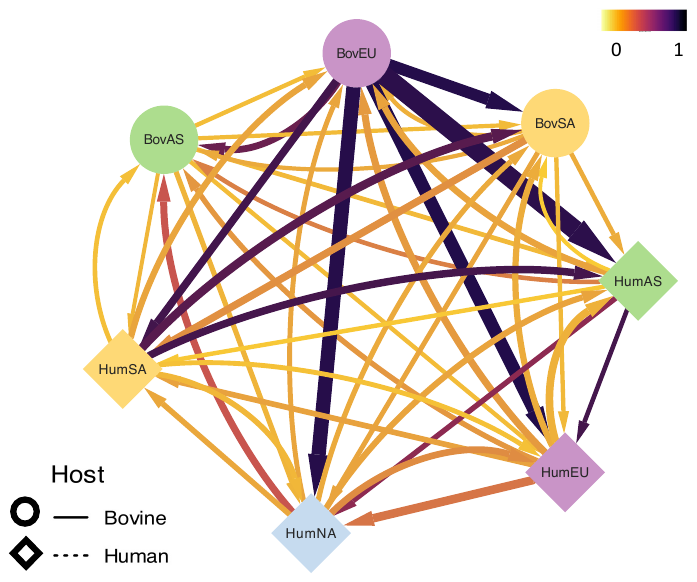

**Fig. S10:** **Network graph of** **BEAST-1 discrete phylogeographic analysis of host/continent stratified subpopulations of Group B Streptococcus clonal group 103/314 genomes (n = 199).** Genomes from Africa are excluded due to an insufficient number of sequences. Arrow thickness scaled to relative migration rate values, arrows coloured according to Bayesian Stochastic Search Variable indicator strength, nodes shaped according to host and coloured by continent. AS Asia, EU Europe, NA North America, SA South America.

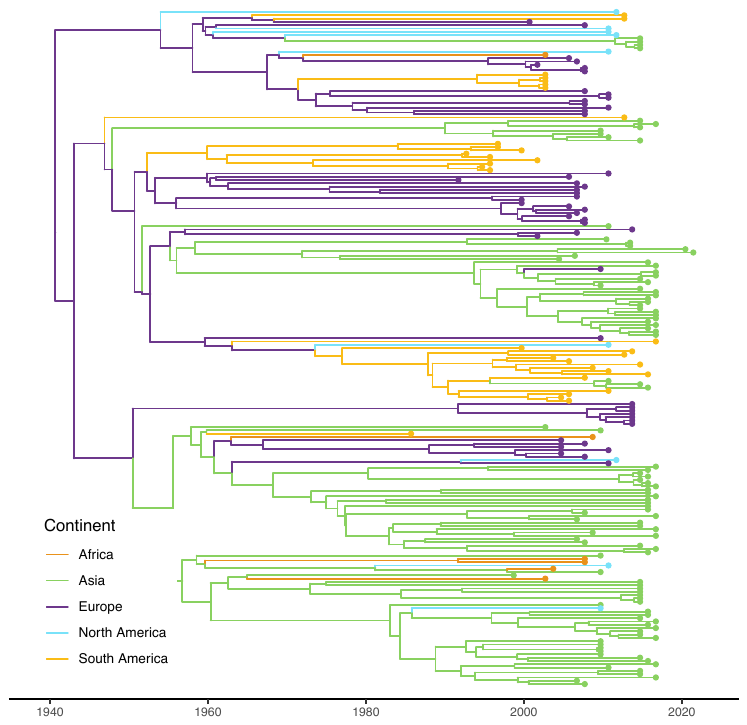

**Fig. S11:** **Time-scaled maximum clade credibility tree based on the alignment of all genomes of Group B Streptococcus clonal groups 103/314 dataset, for which year data were available (n = 245)**. The phylogeny was generated using discrete trait mapping in Bayesian Evolutionary Analysis Sampling Trees (BEAST)-1. The branches are coloured by inferred ancestral continent of origin.
